# Sexual selection does not increase the rate of compensatory adaptation to a mutation influencing a secondary sexual trait in *Drosophila melanogaster*

**DOI:** 10.1101/686741

**Authors:** Christopher H. Chandler, Anna Mammel, Ian Dworkin

## Abstract

Theoretical work predicts that sexual selection can enhance natural selection, increasing the rate of adaptation to new environments and helping purge harmful mutations. While some experiments support these predictions, remarkably little work has addressed the role of sexual selection on compensatory adaptation—populations’ ability to compensate for the costs of deleterious alleles that are already present. We tested whether sexual selection, as well as the degree of standing genetic variation, affect the rate of compensatory evolution via phenotypic suppression in experimental populations of *Drosophila melanogaster*. These populations were fixed for a spontaneous mutation causing mild abnormalities in the male sex comb, a structure important for mating success. We fine-mapped this mutation to an ∼85 kb region on the X chromosome containing three candidate genes, showed that the mutation is deleterious, and that its phenotypic expression and penetrance vary by genetic background. We then performed experimental evolution, including a treatment where opportunity for mate choice was limited by experimentally enforced monogamy. Although evolved populations did show some phenotypic suppression of the morphological abnormalities in the sex comb, the amount of suppression did not depend on the opportunity for sexual selection. Sexual selection, therefore, may not always enhance natural selection; instead, the interaction between these two forces may depend on additional factors.

## Introduction

Sexual selection has important impacts on many aspects of how organisms evolve, including on speciation rates and the degree of sexual dimorphism (e.g., Masta and Maddison 2002; Ellis and Oakley 2016). It was once thought that sexual selection may act independently or even antagonistically to other components of natural selection (e.g. viability and fecundity). However, sexual selection might also be concordant with, and represent a substantial portion of, the total selection on an allele. If so, sexual selection on males might also influence the overall mutation load or rate of adaptation, including in females. For instance, sexual selection may influence how organisms respond to selective pressures in the short term, influencing how quickly populations adapt to novel environments, in particular when the population begins at a distance from an optimum (Long *et al.* 2012). Additionally, some models predict that sexual selection should help populations filter out harmful mutations more rapidly than selection on other fitness components (viability and fecundity selection) alone (Agrawal 2001). This prediction is based partly on the observation that sexual displays are often correlated with overall condition. Any mutation that reduces an organism’s nonsexual fitness is therefore also likely to affect its mating success (Rowe and Houle 1996; Chandler *et al.* 2013b) or even its success in sperm competition (Clark *et al.* 2012). In those cases, total selection against such mutations is stronger than it would be without sexual selection.

Empirical support for this scenario has been mixed. In some studies testing these predictions, evidence supported a role for sexual selection in purging deleterious mutations or accelerating adaptation (Radwan 2004; Sharp and Agrawal 2008; Hollis *et al.* 2009; Jarzebowska and Radwan 2010; McGuigan *et al.* 2011; Long *et al.* 2012; Almbro and Simmons 2014; Lumley *et al.* 2015; Grieshop *et al.* 2016; Jacomb *et al.* 2016). However, a handful of studies also contradict these, perhaps because of the confounding effects of sexual conflict (Hollis and Houle 2011; Arbuthnott and Rundle 2012; Chenoweth *et al.* 2015), or because they used large-effect mutations or strong mutagens not representative of natural variation (Plesnar *et al.* 2011; Cabral and Holland 2014).

Of course, deleterious mutations are not always purged by selection; they can increase in frequency and occasionally become fixed via drift, hitchhiking, or antagonistic pleiotropy, especially if their effects on fitness are only mildly deleterious (and in populations with a small effective population size). In those cases where the deleterious alleles are difficult for selection to purge, alleles at other loci that compensate epistatically for the fitness costs of these fixed deleterious alleles may be favored by selection. There is evidence of compensatory adaptation in both microbial and multicellular organisms. For instance, alleles conferring antibiotic resistance are sometimes costly in the absence of antibiotics, but compensatory mutations can reduce those costs (Reynolds 2000; Maisnier-Patin *et al.* 2002; Comas *et al.* 2012). In the blowfly, diazinon resistance via alleles at the *Rop-1* gene had negative pleiotropic effects, increasing fluctuating asymmetry, but these effects were ultimately compensated by modifiers (McKenzie and Clarke 1988; Davies *et al.* 1996). Additionally, sex chromosome dosage compensation could also be considered a form of compensatory adaptation, having likely evolved in response to loss-of-function mutations on Y or W chromosomes (Charlesworth 1978). In addition, the phenotypic expression (penetrance and expressivity) of many mutations can be strongly influenced by genetic background (e.g., Chandler *et al.* 2013a, 2017; Mullis *et al.* 2018; Hou *et al.* 2019). Thus, selection favoring suppressor alleles at other loci may also contribute to compensatory adaptation by limiting the phenotypic expression of a deleterious mutation.

Although sexual selection has received a lot of attention as a possible influence on the rate of purging of deleterious mutations, the role of sexual selection in compensatory evolution remains largely unexplored. Nevertheless, we might similarly predict that sexual selection can also accelerate compensatory adaptation, especially if sexual displays are condition dependent. In one study (Pischedda and Chippindale 2005), the *nub^1^* mutation, which drastically reduces the size of the wing, resulting in an inhibition of males’ ability to generate courtship songs, was fixed in experimental populations of *Drosophila melanogaster*. As predicted, this mutation had greater fitness costs in males than it did in females, but males also showed greater compensatory fitness recovery over 180 generations (albeit without compensating for the effects on wing morphology directly; A. Chippindale, personal communication), providing some support that sexual selection may enhance compensatory adaptation. However, this study was not replicated (only a single lineage), since the *nub^1^* populations were originally generated for other purposes. Clearly, more study is needed on whether sexual selection can speed up compensatory adaptation.

In this study, we address the question of whether sexual selection can impact the rate of compensatory evolution (via phenotypic suppression) using experimental evolution in *Drosophila melanogaster*. We chose a mutation in the *sex combs distal* gene (*scd^1^*) (Boube *et al.* 1997; Randsholt and Santamaria 2008), a spontaneous, partially penetrant mutation affecting the development of the male sex comb, a structure critical for male mating success (Ng and Kopp 2008) and rapidly evolving across *Drosophila* species (Atallah *et al.* 2009, 2012; Kopp 2011; Malagón *et al.* 2014). First, we further mapped the mutation and characterized its effects across different wild type genetic backgrounds; because we found abundant genetic variation in natural populations modifying its penetrance and expressivity, we next focused on compensatory adaptation via phenotypic suppression in experimentally evolved populations. Despite a general compensatory response, we observed no evidence that sexual selection influenced the rate of compensatory evolution via phenotypic suppression of the sex comb phenotypes.

## Methods

### Mapping scd^1^

*sex combs distal^1^* (*scd^1^*) is a spontaneous X-linked allele resulting in ectopic sex comb bristles on the second tarsal segment of the prothoracic leg in males (Boube *et al.* 1997). In the base stock (Bloomington Drosophila Stock Center strain #5070, *y^1^ scd^1^ ras^1^ v^1^ f^1^*), it has incomplete penetrance, with only about 70% of males showing the ectopic sex comb bristles (Figure 1), and no visible phenotype in homozygous or heterozygous females. In our populations, males also sometimes exhibited minor defects in the primary sex comb, such as a gap or partially untransformed bristles.

**Figure 1.**
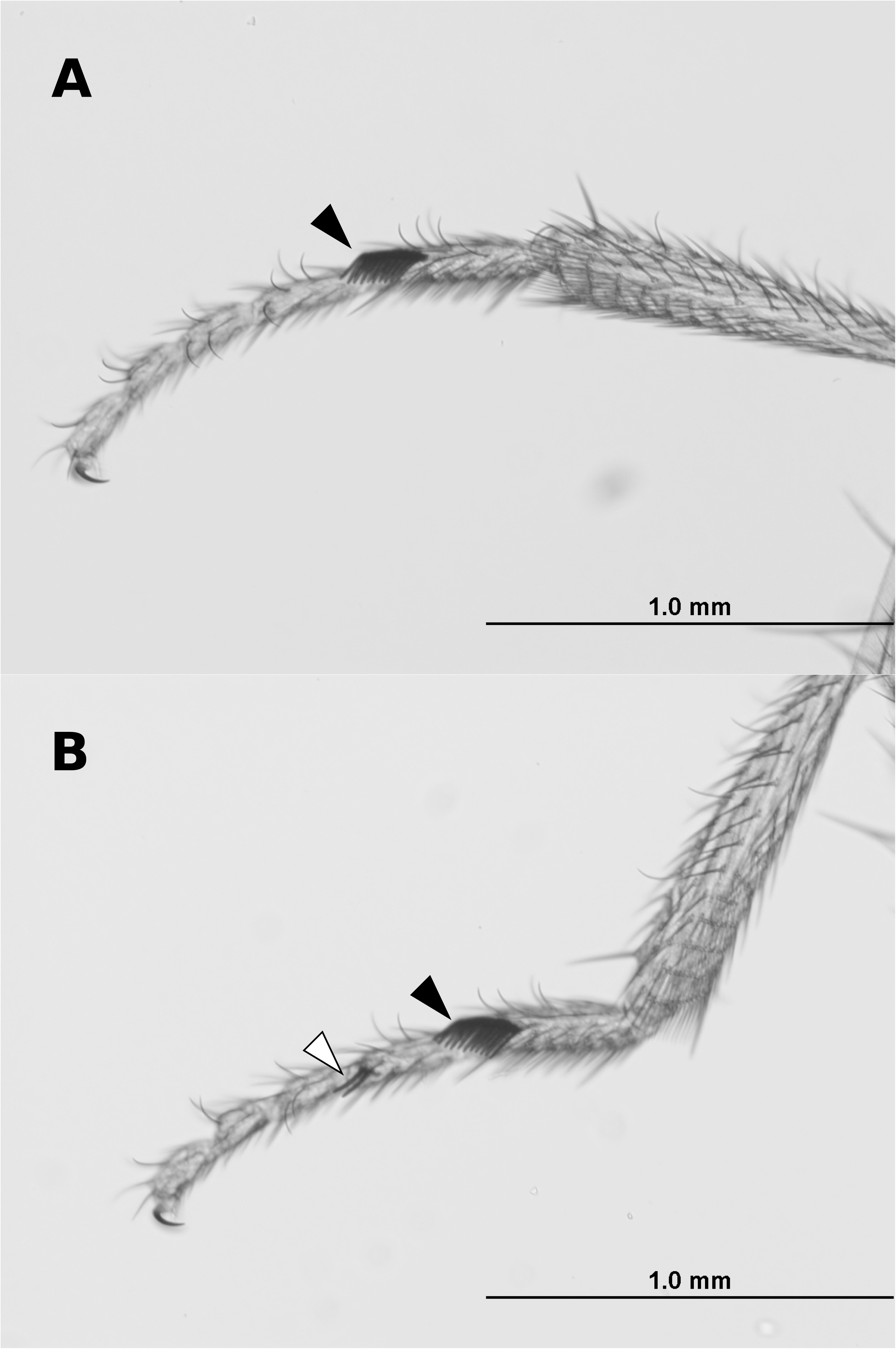
(A) Wild-type *Drosophila melanogaster* leg showing a normal male sex comb (black arrow). (B) *scd^1^* leg showing the normal primary sex comb (black arrow) as well as a smaller ectopic sex comb on the second tarsal segment (white arrow).

The identity of the gene and molecular lesion of this allele are unknown, although some previous recombination mapping suggested it was near 1-30.6, and that a local duplication of the 8C-9B region of the X chromosome onto the Y (*DP(1:Y)FF*), could partially rescue the phenotype of *scd^1^* (Santamaria and Randsholt 1995; Randsholt and Santamaria 2008). We attempted to further fine map the gene through duplication mapping. Virgin female flies of strain BDSC 5070 (*y^1^ scd^1^ ras^1^ v^1^ f^1^*) were crossed to males of strains carrying duplicated segments of the X chromosome translocated onto the Y chromosome or chromosome III (Table 1; Cook *et al.* 2010; Venken *et al.* 2010), and the male offspring were scored for the presence of mutant phenotypes, such as the ectopic sex comb or disruptions in the primary sex comb. Assuming *scd^1^* is a recessive loss-of-function allele, if the duplicated segment contains a functional wild-type copy of the *scd* gene, then the mutant phenotype would be rescued and no male offspring from these crosses would show sex comb defects. Because *scd^1^* is only partially penetrant, we scored at least 20 male offspring from each cross.

**Table 1.**
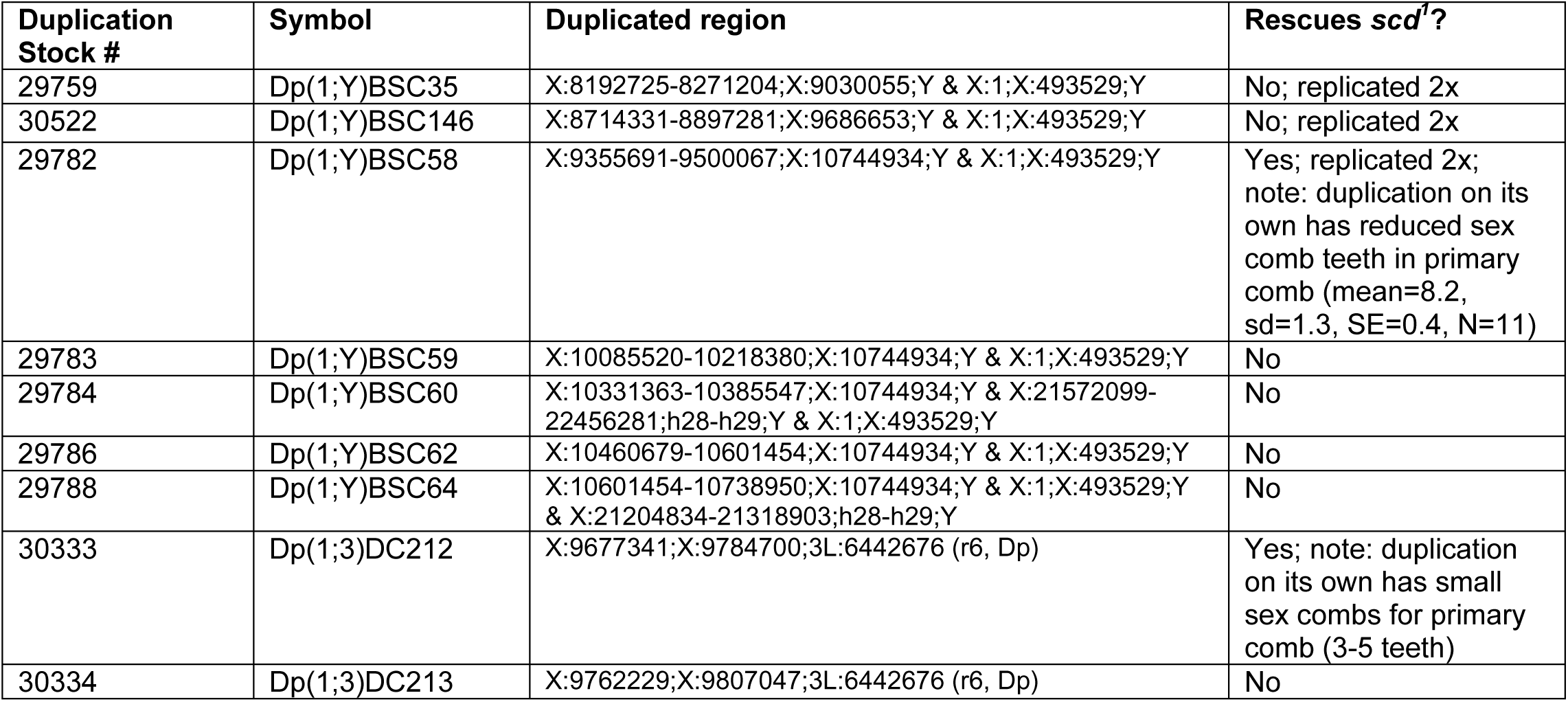
Duplication mapping of the *scd^1^* allele. Homozygous female *scd^1^* flies were crossed to males carrying a segment of the X chromosome duplicated onto either the Y chromosome or chromosome III. Assuming *scd^1^* is a recessive loss-of-function allele, if the male progeny of these crosses show a mutant phenotype, then the duplication does not rescue *scd^1^* and *scd* must lie at least partially outside the duplication. If none of the male offspring show a mutant phenotype, then *scd^1^* is rescued and *scd* lies within the duplication.

### Influence of genetic background on penetrance and expressivity

To determine the extent of genetic variation for the penetrance and expressivity of *scd^1^*, we crossed virgin female *y^1^ scd^1^ ras^1^ v^1^ f^1^* flies to males of a randomly chosen subset of Drosophila Genetic Reference Panel lines (Mackay *et al.* 2012). When the adult F1 offspring eclosed, we fixed specimens in 70% ethanol, and then mounted male prothoracic legs in 70% glycerol and scored them for the presence of ectopic sex combs on the second tarsal segment, abnormalities in the primary sex comb, and primary sex comb tooth number. These crosses only examine each wild-type genetic background in a heterozygous state, and thus it is expected that this will underestimate the actual degree of background dependence, as recessive effects of alleles in each background will not be captured.

To test for an effect of genetic background on penetrance, we fit a logistic model testing for the effect of genotype on presence of an ectopic sex comb using glm() in base R version 3.6.1. We also confirmed those results using glmer() in the lme4 package version 1.1-21.

### Introgression of scd^1^

To generate populations for experimental evolution, we introgressed the *scd^1^* mutation into FVW, a domesticated lab-maintained population founded from flies collected from Fenn Valley, MI in 2010. The FVW population was maintained in population cages with 10 bottles for egg-laying each generation for approximately 10 generations prior to beginning introgressions (Chari *et al.* 2017).

To begin the introgression (Supplementary Figure 1A), virgin females of the 5070 progenitor strain (with the genotype *y^1^ scd^1^ ras^1^ v^1^ f^1^*) were crossed to FVW males; this stock strain carries visible X-linked genetic markers (*y^1^* causes yellow body pigmentation, *ras^1^* and *v^1^* affect eye color, and *f^1^* produces forked bristles). The heterozygous F1 females were then backcrossed to FVW males. From the F2 offspring, we selected males showing the *scd^1^* phenotype, which were thus hemizygous for *scd^1^*, but with phenotypically wild-type eyes and normal bristles to eliminate the *ras^1^*, *v^1^*, and *f^1^* mutations, which are all located to the right of *scd^1^* on the X chromosome. We then crossed these males to virgin FVW females, to obtain female offspring heterozygous for *scd^1^* in a partial FVW background. We then crossed these females to FVW males, and selected males with ectopic sex combs, but not yellow bodies, to eliminate the *y^1^* mutation to the left of *scd^1^*. Each male from these crosses thus carries an independently derived X chromosome with *scd^1^* in a random FVW background.

We obtained eight such males and crossed them to FVW females. From these crosses, we obtained virgin females heterozygous for *scd^1^* in an FVW background. The first of these virgin *scd^1^/scd^+^* females to emerge were crossed with FVW males to obtain more *scd^1^* males with a mostly FVW background. The later-emerging *scd^1^/scd^+^* females were kept isolated at cooler temperatures (18°C) until the *scd^1^* males from the previous cross emerged. We then crossed the *scd^1^/scd^+^* females to the *scd^1^* males. Finally, we set up sib matings among the offspring of these crosses, using only males hemizygous for *scd^1^* (showing a sex comb phenotype) and females of unknown genotype (either *scd^1^/scd^+^* or *scd^1^/scd^1^*). Of those crosses, we kept those in which the mother was inferred to be homozygous for *scd^1^,* in which nearly all male progeny displayed *scd^1^* phenotypes. This allowed us to establish a homozygous *scd^1^* line with an FVW genetic background, which we designated as *scd*\*. *scd*\* carried at least four independently derived X chromosomes with *scd^1^* in an FVW background (Supplementary Figure 1A).

To introduce further genetic diversity (from the FVW population) into *scd*\*, *scd*\* males were crossed to wild-type FVW females to obtain heterozygous *scd^1^* females with additional genetic material from the FVW background. These females were then backcrossed to FVW males. Five replicate backcrosses were set up in culture bottles, each with 25-30 FVW males and 25-30 females, heterozygous for *scd^1^* and for alleles from the FVW background. We then selected males with the *scd^1^* phenotype, and backcrossed them to virgin *scd*\* females, in six replicate bottles each containing 20-25 males and 20-25 females, to maintain *scd^1^* while introducing additional genetic diversity from the FVW population. This whole cycle was then repeated once to establish the *scd*\*\* base population for experimental evolution (Supplementary Figure 1B).

### Fitness effects of scd^1^

To test whether the *scd^1^* allele was deleterious, we tracked changes in the frequency of the *scd^1^* phenotype in polymorphic populations with the *scd^1^* allele at 0.7 initial frequency. We initiated four replicate populations, each consisting of 70 *scd*\*\* males, 70 *scd*\*\* females, 30 FVW males, and 30 FVW females. Populations were placed in population cages with four culture bottles for mating and oviposition for five days, after which the flies were discarded and the bottles transferred to fresh cages at 24°C. After adult flies began emerging, they were allowed to mate for three to four days. The old bottles were then removed, and fresh bottles were placed in the cage for egg laying. After two days of egg laying, the flies were discarded and the bottles moved to fresh cages. This cycle was repeated for a total of nine generations.

For each of the first five generations, and at generation nine, we scored male sex comb phenotypes. 50 males were picked randomly, and the first prothoracic legs from each male were mounted on glass slides in 70% glycerol/PBS to check for the presence of an ectopic second sex comb and other abnormalities. While this does not give an exact measurement of the frequency of the *scd^1^* allele because of this allele’s incomplete penetrance (though penetrance is almost complete in the FVW background; see below), it should provide a reasonable proxy. Even though reductions in the frequency of the mutant phenotype could also be driven by selection for suppressor alleles, this should still give an indication of whether or not the *scd^1^* phenotype is deleterious.

To test whether there was evidence that the *scd^1^* allele was deleterious (and decreased in allele frequency) we fit a logistic regression tracking number of *scd^1^* and wild-type males each generation. As the frequency of *scd^1^* at generation 0 was set at exactly 0.7, we utilized an offset and suppressed the model intercept. Additionally we checked the results of this model using a logistic mixed model allowing for a variation in the slope of the response by replicate lineage. Analyses were conducted in R using glm() and glmer() from the lme4 package.

### Experimental evolution

To test whether sexual selection influences the rate of compensatory adaptation, we set up two treatments. In the low sexual selection (LSS) treatment, we removed sexual selection by enforcing monogamous mating. Each generation, we set up 100 vials, each containing one male and one virgin female. After a three-day interaction period, we anesthetized the flies using CO_2_, discarded males, and placed the females in a population cage with four bottles containing culture media for egg laying. After four days, the bottles were removed and incubated at 24°C. When adult flies began eclosing, we selected virgins for the next generation. Thus, while this treatment did preclude mate choice, it still allowed for fecundity and viability selection (Arbuthnott and Rundle 2012).

In the high sexual selection (HSS) treatment, we followed a similar protocol except allowed the opportunity for sexual selection. Each generation, 100 males and 100 virgin females were allowed to interact in a population cage, along with an open culture bottle for food and moisture. After the three-day interaction period, we placed the cage in a refrigerator to knock the flies out, and then we sorted males and females. Males were discarded, and females were placed in fresh cages with four culture bottles for a four-day egg-laying period. After egg laying, females were discarded, and the bottles were placed in an environmental chamber at 24°C until adults began emerging, at which point we collected virgins for the next generation.

Additionally, we also set up a treatment with low levels of genetic variation (LV) to test whether compensatory adaptation is limited when segregating genetic variation is diminished; in other words, testing whether the mutational target size of the compensatory response was large enough that *de novo* mutations could contribute in the time frame of the experimental evolution regime. In this treatment, each population was established from the offspring of a single-pair mating between a randomly chosen virgin *scd\*\** female and a randomly chosen *scd\*\** male. These populations were kept under the same regime as the HSS treatment.

Finally, we set up a wild-type control (WTC) treatment using wild-type FVW flies. WTC populations were also maintained under the same regime as the HSS treatment. These provide a control for lab domestication and unknown aspects of the experimental protocol.

All populations were initiated using randomly selected *scd*\*\* flies (see above), except for the LV treatments as described. We set up three replicate populations of each treatment except for WTC, in which we performed two replicates. Experimental evolution was conducted for a total of 24 generations. We assayed male sex comb phenotypes as described earlier at generations 1, 7, 13, 19, and 24, using 30 randomly selected males from each population at each time point.

To test how male sex comb traits changed over the course of experimental evolution, we fit generalized linear mixed models using the glmmTMB v0.2.3 (Hadfield 2010) and lme4 v1.1-21 (Bates *et al.* 2015) packages in R (v3.6.1). For sex comb tooth number for both primary and ectopic/secondary combs, we assumed a Poisson distribution and used a log link function. The model included generation, treatment, and their interaction as fixed effects; we also included individual fly, and in some cases replicate population nested within treatment, as random effects (some models failed to converge when replicate nested within treatment was included as a random effect). We also tested for lineage specific zero-inflation in the data, but found no evidence for this, so excluded this to reduce number of parameters. To test whether the frequency of defects in the primary sex comb changed over time, we fit a mixed logistic model (sex comb defects present/absent), again with generation, treatment, and their interaction as fixed effects, and replicate nested within treatment, as well as individual fly, as random effects. For all of these models we examined whether a model with a common intercept (using generation 1 data as the intercept) for each treatment altered estimates relative to models with varying intercepts. In no case did it substantially alter the conclusions, and we present both the common intercept model and varying intercept model in results and supplements. Power simulations were performed using simr v1.0.5 (Green and MacLeod 2016).

### Data availability

All data and scripts are available on Github (https://github.com/DworkinLab/Chandler_etal_G3_2020).

## Results

### Mapping scd^1^

Though we could not map *scd^1^* to a specific gene, we were able to further narrow down its location to an ∼85-kb (cytological region 8F8-9A1) region on the X chromosome through duplication mapping (Table 1, Figure 2). This region contains only two complete annotated protein-coding genes, *btd* and *Sp1* (both of which influence aspects of leg development and morphogenesis), and three annotated long non-coding RNAs, CR42657, CR44016, and CR53498. Interestingly, the two parent strains carrying the duplications that rescued *scd^1^* had smaller than average sex combs in the absence of the *scd^1^* mutation (Table 1), similar to a past study involving this mutation (Randsholt and Santamaria 2008), suggesting that the *scd* gene product is a suppressor of sex comb development.

**Figure 2.**
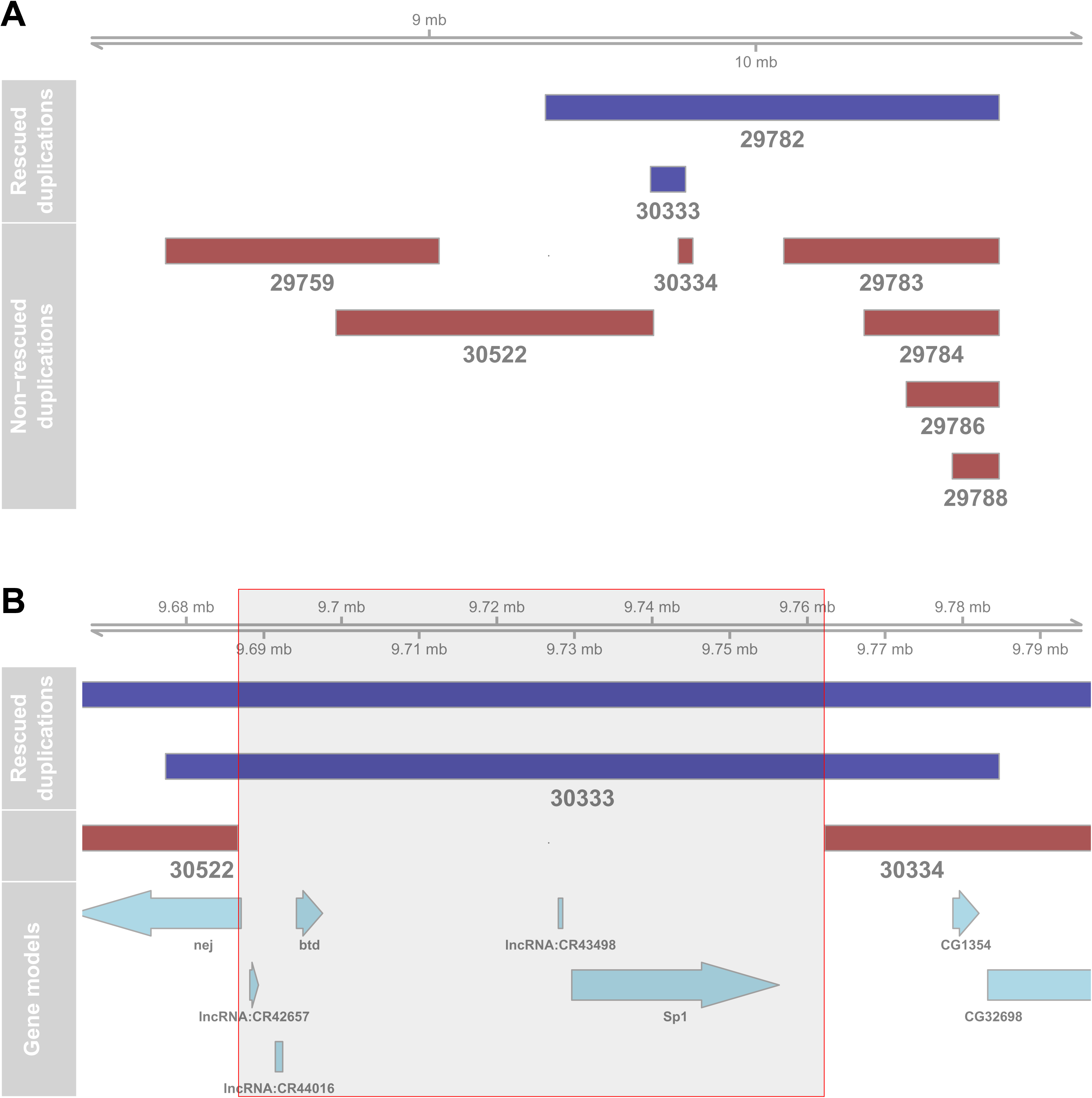
Duplication mapping of *scd^1^*. Purple bars represent duplications that rescued the *scd^1^* mutant phenotype; pink bars represent duplications that failed to rescue *scd^1^*. (A) Entire region of the X chromosome in which duplications were tested. (B) Close-up of the putative *scd^1^*-containing region (red box). Because the duplication carried by strain 30333 is sufficient to rescue the *scd^1^* phenotype, we hypothesize that the *scd* gene must lie entirely within this region; at the same time, the neighboring duplications (30522 and 30334) did not rescue the *scd^1^* phenotype, so *scd* must lie at least partially outside of these regions.

### Influence of genetic background on penetrance and expressivity

When females of the original *scd^1^* stock strain were crossed to males of various wild-type strains to generate males that were hemizygous for *scd^1^* and heterozygous for different genetic backgrounds, the penetrance and expressivity of *scd^1^* varied widely (Figure 3) demonstrating segregating variation for them. A logistic model using penetrance (presence/absence of ectopic sex comb) as the response variable with the genetic background as a fixed effect was a significantly better fit than a null model not accounting for genetic background (x^2^ = 127.6, df = 18, *p* = 5.3 x 10^-19^), and when we fit a model including genetic background as a random effect, there was substantial among-strain variance (σ^2^ = 4.19 on the link scale). Similarly, a model with number of ectopic sex comb teeth as the response variable (expressivity) and genetic background as a fixed effect was a significantly better fit than the null model (x^2^ = 259.0, df = 18, *p* < 1.0 x 10^-10^). The progenitor *scd^1^* strain from the Bloomington stock center had the lowest penetrance (frequency of flies exhibiting an ectopic sex comb on the second tarsal segment) and among the lowest expressivity (number of teeth in the ectopic sex comb). Some of the other wild-type genetic backgrounds, even in a heterozygous state, resulted in nearly complete penetrance for *scd^1^*, including the outbred population (FVW) used for experimental evolution (described below). Interestingly, the FVW outbred population only showed intermediate levels of expressivity of the mutant phenotype, consistent with segregating variation in this population. Overall this result suggests genetic background has a strong impact on the phenotypic expression on *scd^1^*. It also suggests that the partial penetrance initially observed in the progenitor stock (strain 5070) may reflect the accumulation of suppressor/compensatory mutations in the base stock center strain itself. These results suggest that a compensatory response could potentially be due to the accumulation of segregating suppressor alleles.

**Figure 3.**
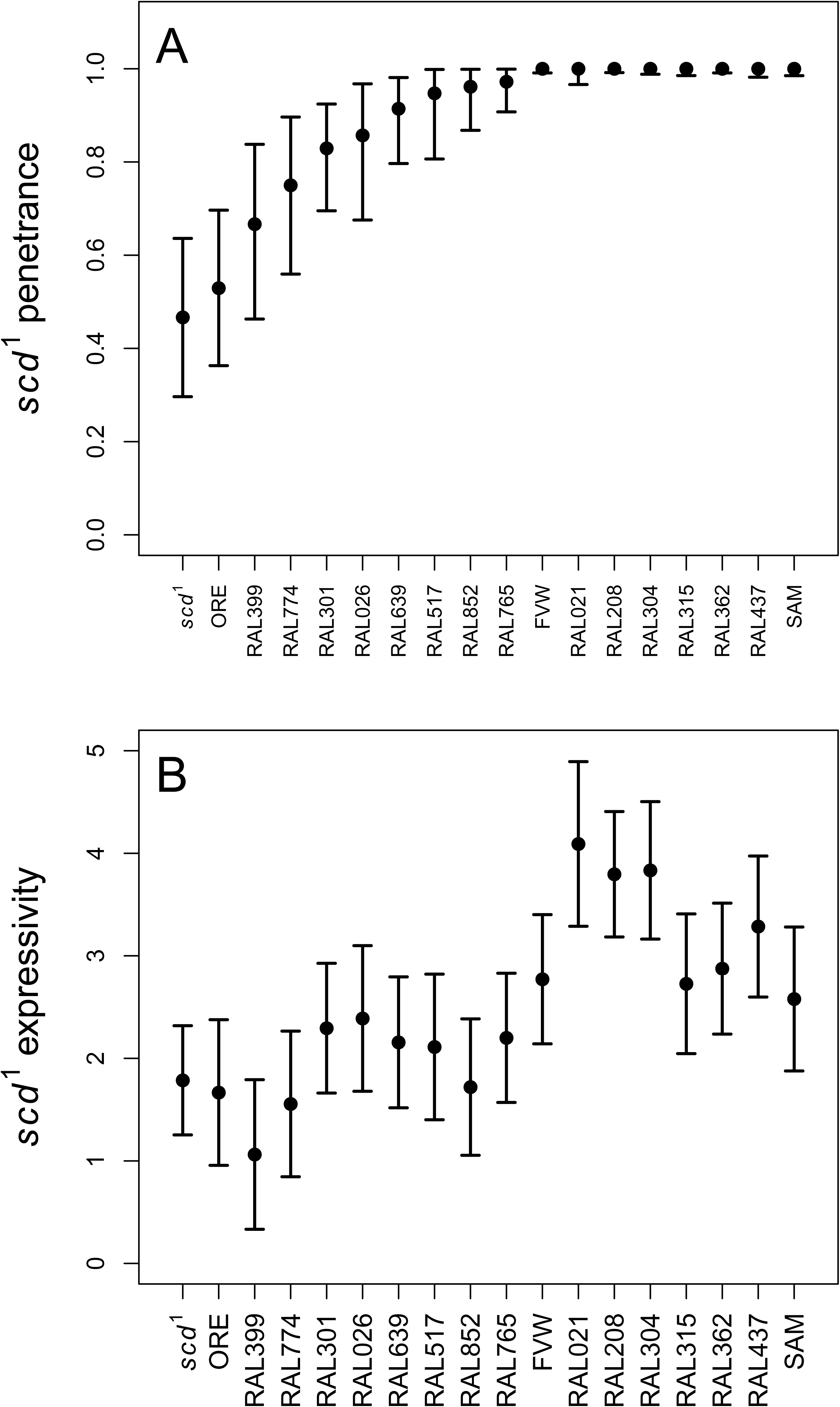
(A) Penetrance and (B) expressivity of the *scd^1^* mutation varies among different wild type genetic backgrounds. In these experiments, stock *scd^1^* females were crossed to males of different wild-type strains, and phenotypes were scored in F1 males (which were hemizygous for *scd^1^* and heterozygous for a different wild genetic background). Penetrance was measured as the proportion of males showing an ectopic sex comb on the second tarsal segment of the prothoracic leg, while expressivity was measured as the number of sex comb teeth on the ectopic sex comb. Error bars indicate 95% confidence intervals.

### Fitness effects of scd^1^

The frequency of male flies exhibiting the *scd^1^* phenotype (in an FVW genetic background) decreased across five generations of experimental evolution in populations polymorphic for *scd^1^* (Figure 4). With a starting allele frequency of 0.7 the frequency decreased to an average frequency of 0.4 (across the multiple replicates) in males by generation 9. To test this more rigorously, we fit a logistic model with an offset (starting frequency of *scd*^1^ = 0.7), and the effect of generation was significant (effect = −0.197 on logit link scale, s.e. = 0.049, *p* = 5.1 x 10^-5^). These results are consistent with the mutation having moderate deleterious effects.

**Figure 4.**
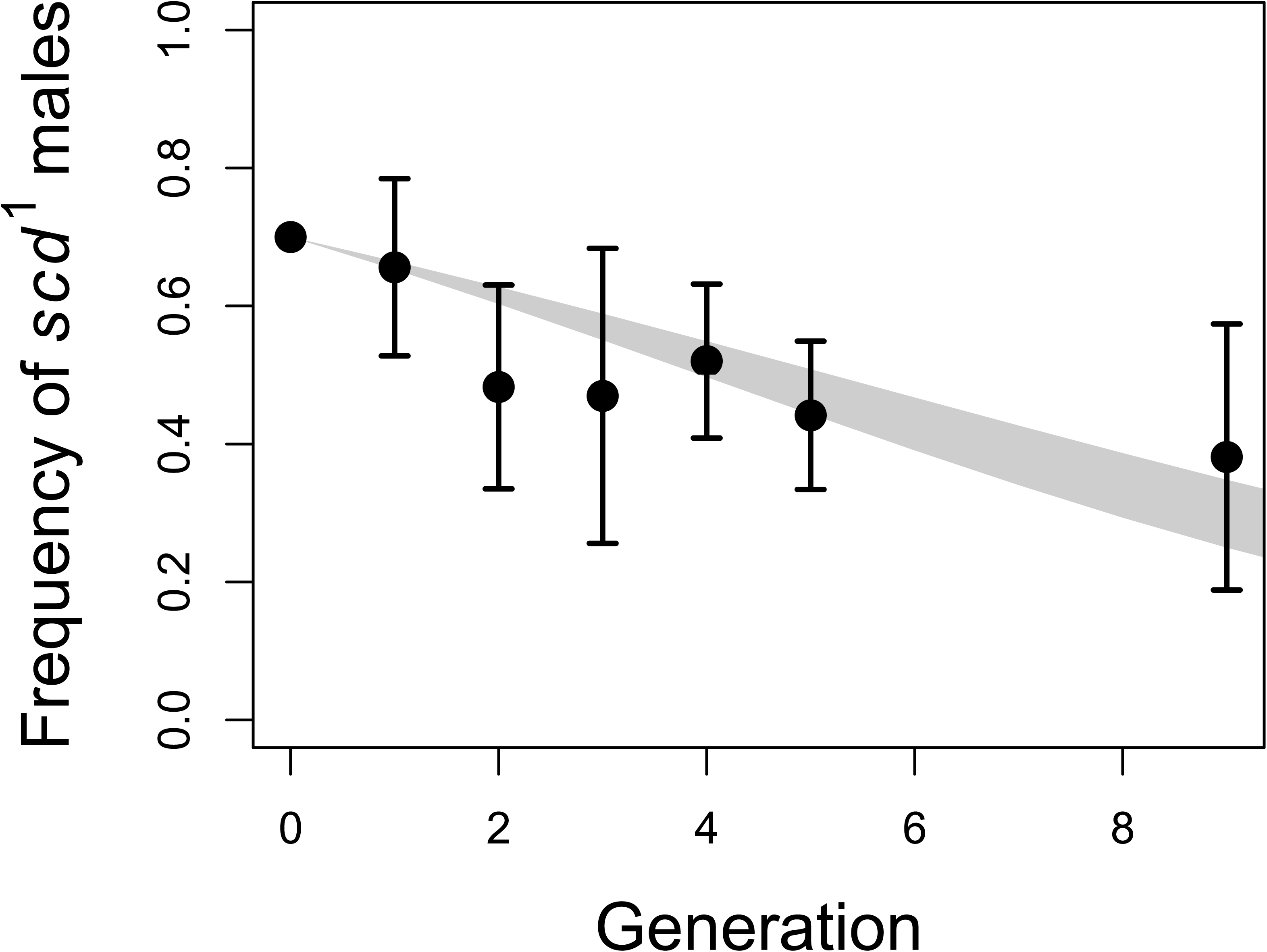
The *scd^1^* mutation is deleterious. Average frequency of *scd^1^* over time, across 4 replicate populations, each initialized with 70% *scd^1^* males. Error bars indicate 95% confidence intervals. The shaded region indicates the 95% confidence interval for the best-fit line, with the starting frequency fixed at 70%.

### Experimental evolution

In the populations carrying the *scd^1^* allele, defects such as gaps in the primary sex comb were observed occasionally, and at significantly higher frequencies in the High Sexual Selection (HSS) populations, and marginally significant frequencies in the Low Sexual Selection (LSS) populations, than in the wild-type populations. The frequency of these gaps appeared to decrease in the HSS populations across the 24 generations of the experiment, although the interaction between generation and treatment was not significant (Figure 5B; Table 2). In addition, the ectopic sex combs induced by the *scd^1^* mutation became smaller on average in both the HSS and LSS treatments (Figure 5C; Table 3), losing on average ∼0.5 teeth across the 24 generations of experimental evolution, though the effect of generation was only marginally significant in the HSS treatment. This is consistent with some of the compensatory response being the result of the increase in frequency of naturally occurring suppressor alleles. When we allowed the intercepts to vary for the HSS and LSS treatments, results were similar, with a significant main effect for generation, but no significant generation-by-treatment interaction (Supplementary Tables 1 and 2). Thus we saw no evidence for differences in rate of compensation between the HSS and LSS treatments, with the magnitude of the interaction term (change in slope relative to LSS) being much smaller than the effect of generation (i.e. overall compensatory response). Indeed, the slope of the compensatory response is weaker for the HSS treatment (relative to LSS), counter to our predictions. This suggests any additional compensatory effects of sexual selection (i.e. after accounting for viability and fecundity selection) were relatively weak in this experimental system. Using a power analysis, we confirmed that the power to detect such an effect would be very small (Supplementary Figure 2) unless we used a large number of independent replicate lineages (∼30 per treatment), although the power to detect an effect of this magnitude (assuming it was real) would be approximately 80% with three replicates if the response continued for 35 or more generations of experimental evolution (Supplementary Figure 3). As a thought experiment (as this variable is not under experimental control, but was being estimated), we performed a power analysis where we modified the magnitude of the slopes of compensatory response (between LSS and HSS). Assuming the estimated effect is real (just with high degree of uncertainty due to sampling) the power for our experimental design would be ∼18% (95% CIs 11 - 26.9%). If the difference between the LSS and HSS treatments is of similar magnitude to that of the overall compensatory response the power increases to ∼34% (CIs 24.8 – 44.1%), and reaches power of 76% (CIs 66.4 – 84.0 %) when the magnitude of the difference between LSS and HSS is 100% greater than the observed slope.

**Figure 5.**
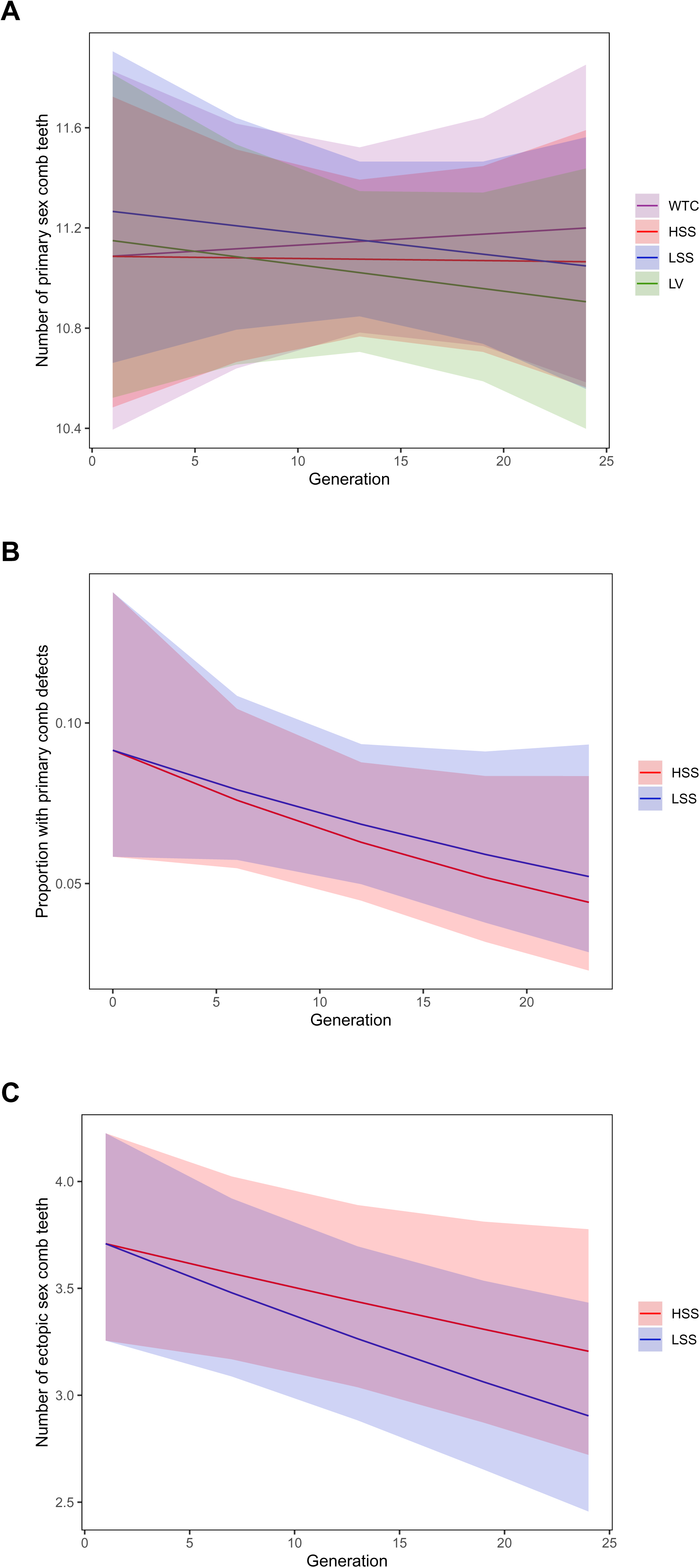
Plots of model fits for sex comb traits in experimental evolution populations. (A) Number of teeth in the primary sex comb across 24 generations of experimental evolution in all experimental treatments. (B) Proportion of male flies with defects in the primary sex comb in the HSS and LSS treatments; we fit a model with a shared intercept for these two treatments. (C) Number of teeth in ectopic/secondary sex combs in males in the HSS and LSS treatments, again with a common intercept. Shaded regions on plots indicate 95% confidence intervals for the predicted values.

**Table 2.** Effect estimates for logistic mixed model for gaps in the primary sex comb as response variable. This model had included only the HSS and LSS treatments, and these two treatments were constrained to have a common intercept, but not slope. The confidence intervals for the effect of generation in each treatment overlap.

|  | Estimate | Standard Error | *P* |
| --- | --- | --- | --- |
| Intercept | -2.30 | 0.25 | $< 2.0 \times 10^{-16}$ |
| generation-HSS | -0.034 | 0.021 | 0.106 |
| generation-LSS | -0.026 | 0.020 | 0.188 |

**Table 3.** Effect estimates for generalized linear model (poisson) using number of teeth in the ectopic/secondary sex comb as response variable. This model included only the HSS and LSS populations; WTC populations were excluded because they did not display ectopic sex combs, and because the purpose of this model was to test for a difference between the HSS and LSS treatments specifically, we considered the LV treatment on its own separately. The HSS and LSS treatments were constrained to have a common intercept in this model, but were allowed to have different slopes. The confidence intervals for the effect of generation in each treatment overlap.

|  | Estimate | Standard Error | *P* |
| --- | --- | --- | --- |
| Intercept | 1.31 | 0.066 | $< 2.0 \times 10^{-16}$ |
| generation-HSS | $-6.3 \times 10^{-3}$ | $3.7 \times 10^{-3}$ | 0.085 |
| generation-LSS | -0.011 | $3.8 \times 10^{-3}$ | 0.005 |

For the Low genetic Variation (LV) treatment no significant change in number of ectopic sex comb teeth over time was observed, as expected (generation effect = 0.0017, SD = 0.0044, *p* = 0.70). No significant changes were observed in the primary sex comb tooth number across 24 generations in any of the experimental treatments (Figure 5A; Table 4).

**Table 4.** Effect estimates for generalized linear mixed model for primary sex comb tooth number as response variable; generation, treatment, and their interaction as fixed effects; and individual fly as a random effect (models including replicate population nested within treatment failed to converge).

|  | Estimate | Standard Error | *P* |
| --- | --- | --- | --- |
| Intercept | 2.41 | 0.035 | $< 2 \times 10^{-16}$ |
| generation | $4.4 \times 10^{-4}$ | $2.2 \times 10^{-3}$ | 0.85 |
| treatment-HSS | $4.2 \times 10^{-4}$ | 0.046 | 0.99 |
| treatment-LSS | 0.017 | 0.045 | 0.71 |
| treatment-LV | $6.9 \times 10^{-3}$ | 0.047 | 0.88 |
| generation x treatment-HSS | $5.2 \times 10^{-4}$ | $2.9 \times 10^{-3}$ | 0.86 |
| generation x treatment-LSS | $1.3 \times 10^{-3}$ | $2.9 \times 10^{-3}$ | 0.66 |
| generation x treatment-LV | $1.4 \times 10^{-3}$ | $3.0 \times 10^{-3}$ | 0.64 |

## Discussion

Some, but not all, previous work has found that sexual selection may facilitate populations in purging deleterious mutations (Radwan 2004; Hollis *et al.* 2009; Jarzebowska and Radwan 2010) or accelerating rate of adaptation (Jacomb *et al.* 2016; Parrett and Knell 2018). Few studies, however, have addressed whether sexual selection may facilitate compensatory adaptation, where populations evolve traits to compensate for the fitness costs of deleterious mutations. While compensatory adaptation itself is well documented in other systems (Reynolds 2000; Maisnier-Patin *et al.* 2002; Estes *et al.* 2011; Chandler *et al.* 2012; Comas *et al.* 2012; Chari *et al.* 2017), in this experiment, we found no effect of the sexual selection regime on the rate of compensatory adaptation (at least with respect to the mutation’s sex comb phenotypes). This is perhaps surprising, because we found clear evidence of standing genetic variation modulating the expression of this mutation, so genetic variation does not appear to be a limiting factor here. Moreover, our experiments show that the *scd^1^* mutation is deleterious (Figure 4), and given the importance of the *Drosophila* sex comb for male mating success (Ng and Kopp 2008), we expected that the fitness costs of this mutation would involve male sexual fitness. Thus, we predicted that the costs of this mutation would be higher in the HSS treatment, in which there was a high opportunity for female mate choice, than in the LSS treatment, with reduced opportunity for sexual selection. It is possible that this mutation has effects on other aspects of fitness in males or females (viability, fecundity), but unfortunately our experiments did not directly measure specific fitness components. Even so, theoretical work predicts that sexual selection should act in concert with natural selection because of condition dependence (Whitlock and Agrawal 2009); that is, mutations that reduce nonsexual fitness should also reduce mating success, since sexual displays are often indicators of overall condition. While some work has supported this prediction, our findings add to a growing body of work suggesting that this is not always the case (Hollis and Houle 2011; Plesnar *et al.* 2011; Arbuthnott and Rundle 2012, 2014; Cabral and Holland 2014; Power and Holman 2015; Chenoweth *et al.* 2015). It is possible that the relatively minor degree of phenotypic suppression observed here made it difficult to detect differences between treatments.

Even though sexual selection did not impact the rate of compensatory evolution, we did observe evidence of weak compensatory adaptation via phenotypic suppression of the *scd^1^* mutation in both the HSS and LSS treatments. On average, the ectopic sex combs lost about half a tooth (starting with a mean of ∼ 3.5 teeth) over the course of 24 generations in these populations; in other words, the expressivity of the mutation declined slightly. One possible explanation for the similar response in both the high and low sexual selection treatment is simply that the compensatory response (in terms of phenotypic suppression) was sufficiently weak that any subtle difference between these treatments would be difficult to detect given our design. However, the power analyses (Supplementary Figures 2 and 3) suggest that if an effect of this magnitude were real, it is sufficiently small that it would require ∼25 replicate lineages of each treatment to detect or a doubling of the number of generations of experimental evolution.

As expected, we did not observe any significant trend in the LV treatment, in which populations experienced genetic bottlenecks prior to beginning the experiment (LV populations were treated the same way as HSS populations). Combined with the observation that genetic background has strong influences on the penetrance and expressivity of this mutation (Figure 3), this suggests that compensatory adaptation by phenotypic suppression relies heavily on the presence of standing genetic variation, rather than rapid accumulation of new mutations. An interesting side note is that the initial strain (obtained from the Drosophila stock center) carrying *scd^1^* has among the lowest penetrance/expressivity for this mutation of all the genetic backgrounds that we tested. This may suggest that the stock strain has already undergone compensatory adaptation, and that alleles suppressing the phenotypic expression of *scd^1^* had become fixed throughout the maintenance of this stock (which were subsequently removed when we outcrossed the mutation), though unfortunately we do not have any data on how long the *scd^1^* stock strain has been maintained.

While we were unable to map *scd^1^* to a specific gene, we were able to localize it to an ∼85 kb region containing only a few candidates. The only protein-coding candidate genes in this region, *Sp1* and *btd*, both have known roles in leg development (Estella and Mann 2010), but are not specifically known to influence sex comb development. There are also three long non-coding RNAs in this region (CR42657, CR44016, and CR43498). Interestingly, all three of these RNAs show evidence of male-specific expression in modENCODE RNA-seq data available on FlyBase (Graveley *et al.* 2011; Brown *et al.* 2014). However, CR44016 shows expression only at very low levels and only in adult males, not pupae or larvae, suggesting it is unlikely to be involved in the development of sex combs. CR42657 and CR43498 both show expression in pupae and/or larvae, as well as adult males (but not adult females); however, these RNAs seem to be expressed in the testis and accessory gland and not other tissues (though expression in legs specifically was not assessed in the modENCODE dataset). This suggests that these male-specific reproductive tissues may be driving these sex-specific expression patterns, not a role in sex comb development. Further work is necessary to identify the molecular nature of *scd^1^*.

There are a number of important limitations to point out about our study. First, much of the focus was on compensation by suppression of the phenotypic effects of the *scd^1^* mutation on the sex combs directly. While we observed similar levels of phenotypic compensation with both our high and low sexual selection treatments (LSS and HSS), it is possible that compensatory evolution differed with respect to the fitness components (viability, fecundity, and sexual/mating components), which were not evaluated. Thus we limit our interpretation to the effects on morphological compensation/suppression, recognizing that we cannot rule out differential patterns of compensatory response for fitness *per se*. Indeed, this pattern has been observed previously (Pischedda and Chippindale 2005; Chari *et al.* 2017). Additionally, this experiment was performed over a relatively short time period (25 generations); if we continued the experiment over a longer period, subtle differences in the rate of morphological compensation may have become apparent, as indicated by our power analysis (Supplementary Figure 3).

In summary, we found evidence of moderate compensatory adaptation to a deleterious mutation by selection for modifier alleles that suppress the mutation’s phenotypic effects. However, while compensatory adaptation did depend on the presence of standing genetic variation, increased opportunity for sexual selection did not increase the rate of compensatory response, in spite of the affected phenotype’s known role in mating. Indeed, the HSS treatment showed a slightly weaker (but not significantly different) rate of compensatory response. However, as shown in the power analysis, additional replicates, or following these lineages for a longer evolutionary trajectory would be helpful to reduce the uncertainty due to sampling. Nonetheless, this adds to a growing body of studies suggesting that sexual selection does not always enhance natural selection. Future work should tease apart when and why sexual and natural selection act in concert and when they are likely to operate differently (Martínez-Ruiz and Knell 2017).

## Acknowledgments

We thank Shelby Hemker and Cody Porter for assistance in data collection and fly maintenance, as well as the editor and two anonymous reviewers for their feedback on earlier versions of this manuscript. The BEACON Center for the Study of Evolution in Action (NSF DBI-0939454) provided support for this work. ID was supported by funds from NSERC (Canada) Discovery and Discovery Accelerator awards.

---

**Supplementary Table 1.** Effect estimates for logistic mixed model for gaps in the primary sex comb as response variable; generation, treatment, and their interaction as fixed effects; and replicate nested within treatment, and individual fly as random effects. This model included only the HSS and LSS treatments, as the WTC treatment was not expected to have appreciable frequencies of sex comb defects, and the LV treatment was considered separately.

|  | Estimate | Standard Error | *P* |
| --- | --- | --- | --- |
| Intercept | -1.98 | 0.33 | 1.4 x 10<sup>-9</sup> |
| generation | -0.054 | 0.026 | 0.037 |
| treatment-LSS | -0.66 | 0.50 | 0.19 |
| generation x treatment-LSS | 0.048 | 0.036 | 0.18 |

**Supplementary Table 2.** Effect estimates for logistic mixed model for number of teeth in the ectopic sex comb as response variable; generation, treatment, and their interaction as fixed effects; and replicate nested within treatment, and individual fly as random effects. This model included only the HSS and LSS treatments, as the WTC treatment was not expected to have ectopic sex combs, and the LV treatment was considered separately.

|  | Estimate | Standard Error | *P* |
| --- | --- | --- | --- |
| Intercept | 1.37 | 0.089 | $< 2 \times 10^{-16}$ |
| generation | -0.0077 | 0.0038 | 0.0489 |
| treatment-LSS | -0.12 | 0.13 | 0.34 |
| generation x treatment-LSS | $-1.5 \times 10^{-3}$ | $5.6 \times 10^{-3}$ | 0.79 |

**Supplementary Figure 1.**
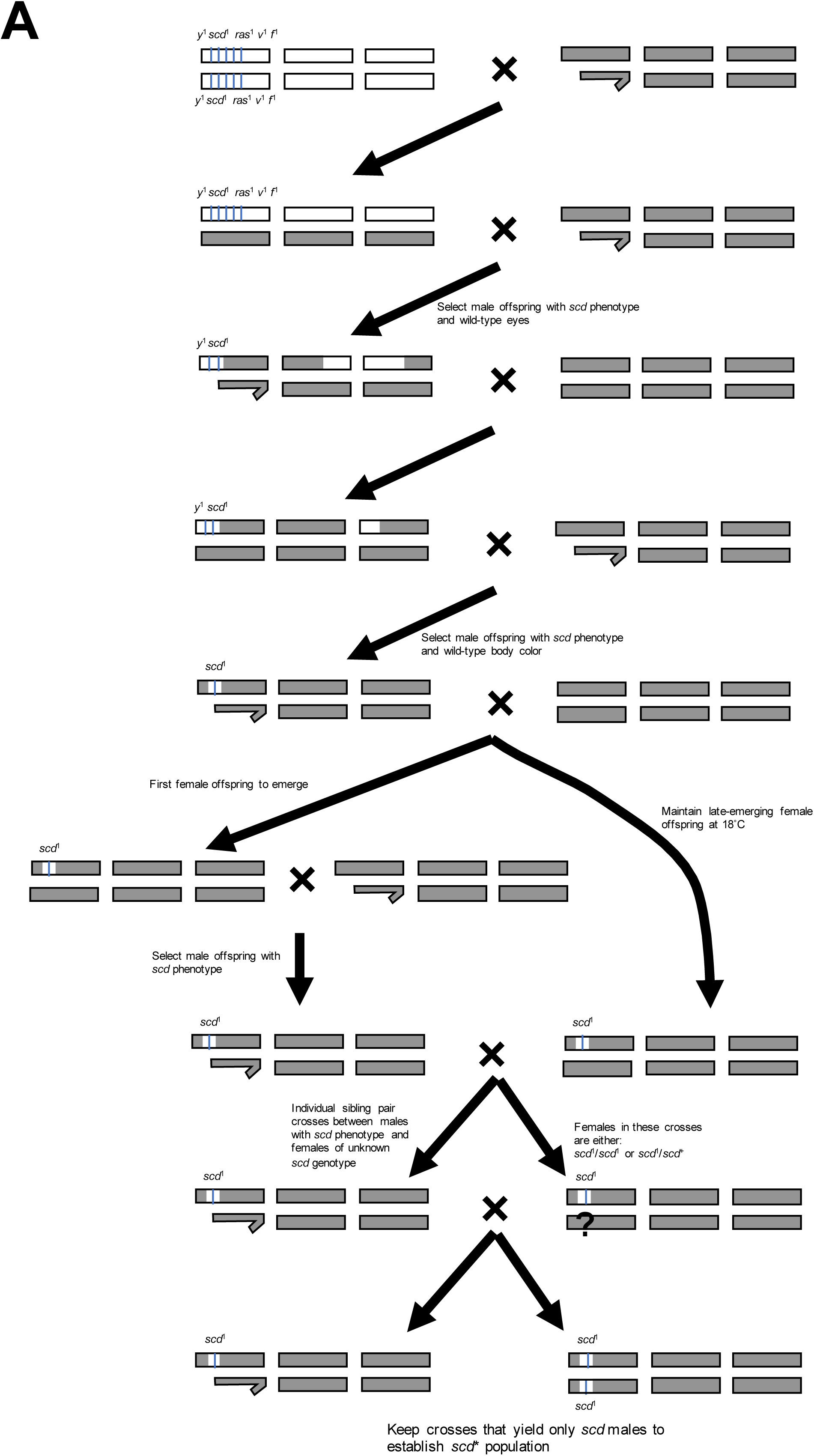
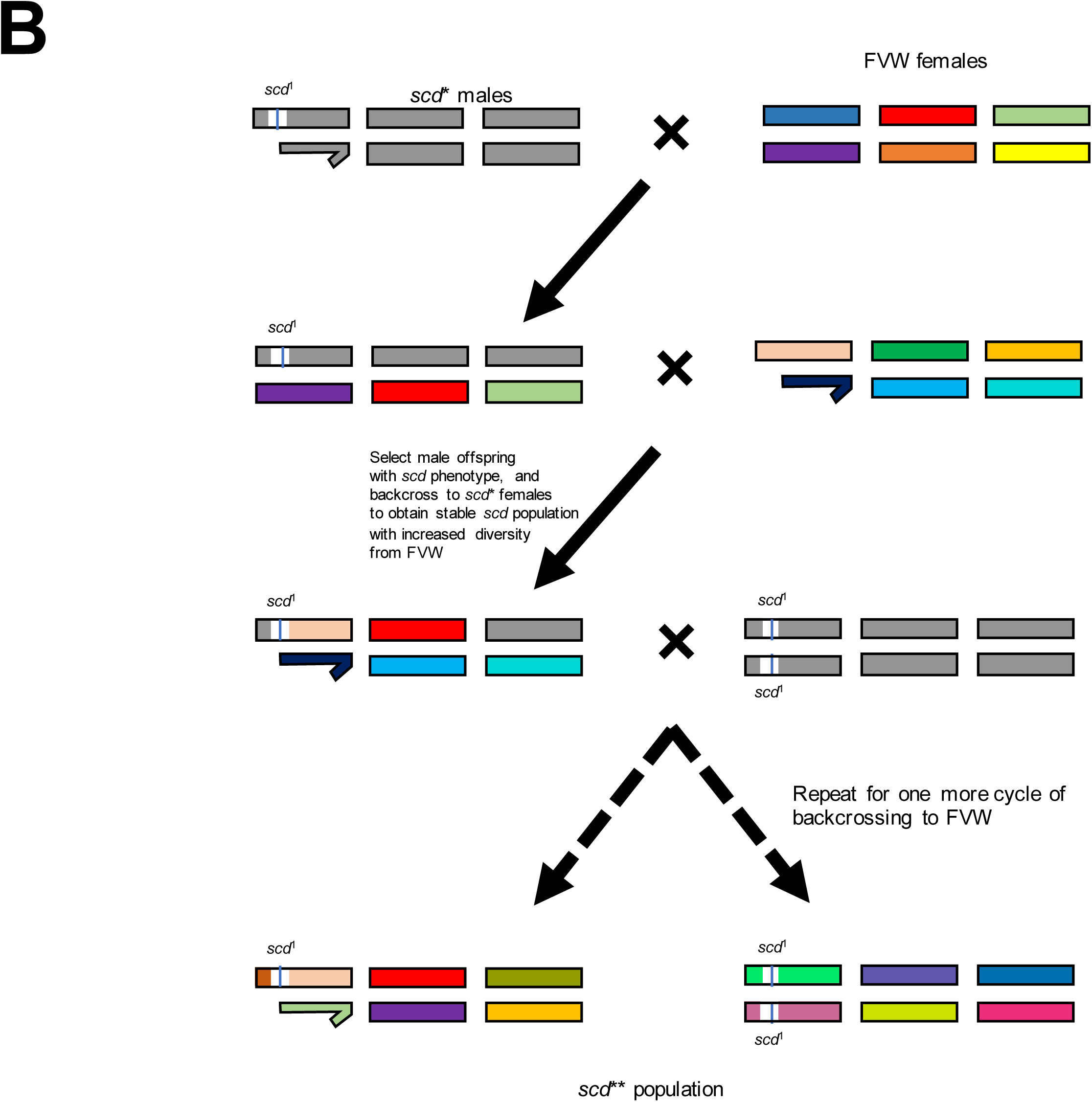
Crossing scheme for introgression of *scd*^1^ into the FVW wild-type genetic background. (A) Initial introgression of *scd*^1^ into FVW to establish the *scd*\* population. (B) Further backcrossing using larger numbers of flies to introduce additional genetic diversity from FVW to establish the *scd*\*\* base population for experimental evolution. See Methods section for detailed description of introgression scheme.

**Supplementary Figure 2.**
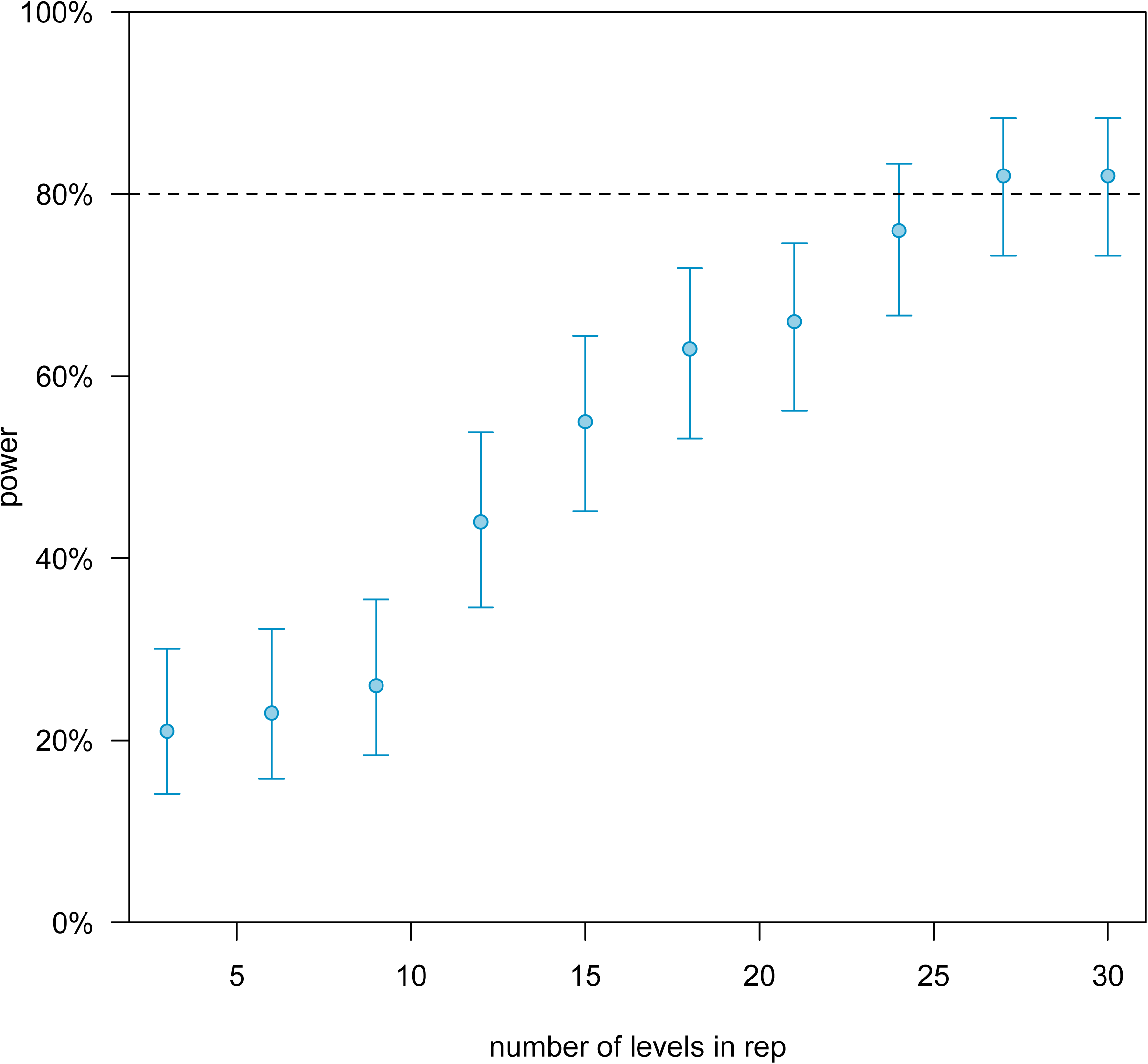
Statistical power to detect a significant effect of sexual selection regime on compensatory adaptation as a function of number of replicate lineages in each treatment, holding all other aspects of the experiment constant. This analysis assumes the magnitude of the effect of sexual selection regime is similar to what we actually observed. At least 25 replicate lineages are necessary to detect such an effect 80% of the time.

**Supplementary Figure 3.**
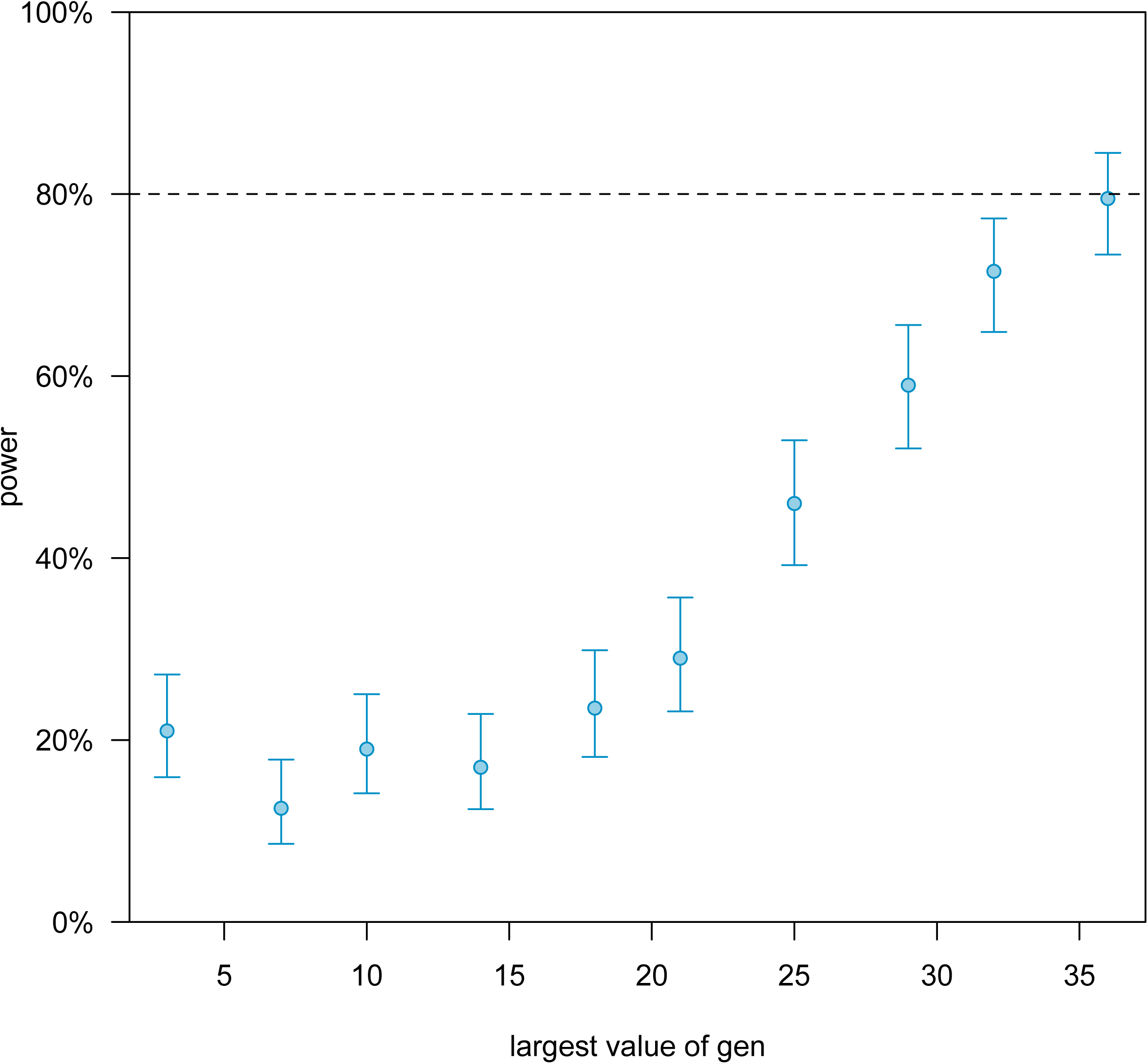
Statistical power to detect a significant effect of sexual selection regime on compensatory adaptation as a function of number of generations of experimental evolution, holding all other aspects of the experiment constant. This analysis assumes the magnitude of the effect of sexual selection regime is similar to what we actually observed. Approximately 35 generations are necessary to detect such an effect 80% of the time.

